# New three-dimensional preclinical models to understand and treat liver cancers activated for the β-catenin pathway

**DOI:** 10.64898/2026.04.01.715868

**Authors:** Vanessa Bou Malham, Fanny Léandre, Akila Hamimi, Isabelle Lagoutte, Sandrine Bouchet, Angélique Gougelet, Sabine Colnot, Christèle Desbois-Mouthon

## Abstract

**Background & aims:** Constitutive activation of the β-catenin pathway is a determining feature in the pathogenesis of two primary liver cancers, namely HCC and hepatoblastoma (HB). Activating alterations in *CTNNB1* gene and, to a lesser extent, inhibiting alterations in *APC* gene are observed in 30 to 40% of HCC cases and 80 to 90% of HB cases. For both tumours, therapeutic management is far from optimal. Therefore, relevant experimental models are needed to increase our knowledge and test new therapeutic approaches.

**Methods:** Organoids and tumouroids were established from APC^Δhep^ and βcat^Δex3^ mouse models, which are clinically relevant models for β-catenin-activated HCC and mesenchymal HB. We developed a new methodological approach based on a dynamic suspension culture in a rotating bioreactor. Morphological and molecular characteristics and sensitivity to WNTinib, a treatment already successfully tested on human HCC and HB tumouroids, were evaluated by histology, immunohistochemistry, immunofluorescence, and RT-qPCR.

**Results:** This easy-to-implement methodology allows for the rapid generation of a large number of organoids and tumouroids that are uniform in size and show no signs of cell death in their core. The robustness of the methodology is illustrated by the maintenance of the histological architecture, cell diversity and gene expression in organoids and tumouroids in comparison with the native liver tissues. In addition, the value of the HCC-derived tumouroids for evaluating cancer treatment was assessed based on their responsiveness to the β-catenin antagonist WNTinib.

**Conclusions:** The organoids and tumouroids that we present here are new reliable *in vitro* cancer models, recapitulating the main features of β-catenin-driven HCC and mesenchymal HB. They can be integrated into an appropriate platform for drug screening and could enable the development of “à la carte” therapies that are urgently needed for these indications.

**Impact and implications:** This study addresses the critical need for representative *in vitro* models to investigate β-catenin-driven liver cancers. The organoids and tumouroids developed here are particularly valuable for researchers seeking robust, reproducible models that accurately reflect the cellular diversity and gene expression profiles of native liver tumours. These findings have practical applications in exploring cancer mechanisms, screening new drugs, optimizing personalized treatment strategies, and reducing reliance on animal models, which ultimately benefits patients.

**Highlights:**

- Easy and rapid generation of mouse liver organoids and tumouroids from β-catenin activated tumours using culture in a bioreactor
- Tumouroids preserve histology, cell diversity, and gene expression of native tissue
- HCC-derived tumouroids respond to β-catenin inhibitor WNTinib
- These reliable 3D models reduce reliance on animal experiments for drug testing

## Introduction

Oncogenic activation of β-catenin is a driving event in the development of primary liver cancers in adults and children, namely HCC and hepatoblastoma (HB), respectively. HCC is the most common primary liver tumour with > 900,000 newly diagnosed people per year in the world. Its incidence is likely to increase in the coming years, making it a major public health problem. HCC also stands at the third rank of death-related to cancer worldwide accounting for more than >830,000 deaths in 2020 (1). HB is a much rarer liver tumour but is the most frequent childhood liver cancer, developing before five years of age. HB represents 1-1.5 cases per million children and its incidence has tripled over the last 30 years (2).

In these primary liver cancers, recurrent mutations and deletions affecting *CTNNB1*, the gene encoding β-catenin, have been reported. Gain-of-function mutations in *CTNNB1* are found in 30 to 40% of HCC while gain-of-function deletions in *CTNNB1* are found in 80% of HB. Loss-of-function deletions in *APC,* a tumour-suppressor gene encoding the main negative regulator of the β-catenin pathway, have been also reported in about 3 and 10% of HCC and HB, respectively (3, 4). These alterations lead to β-catenin stabilization and accumulation in the nucleus, where it binds to the transcription factor TCF4 and regulates gene expression to promote tumour initiation and progression. β-catenin pathway-mutated HCC represents a homogenous subgroup of differentiated and low proliferating tumours whose prognosis may be better than other subtypes if diagnosed at an early stage and resected (2). Unfortunately, there is no specific treatment for advanced *CTNNB1*-mutated HCC, and it appears that oncogenic activation of β-catenin confers frequent innate resistance to the reference treatment combining immune checkpoint inhibitor plus antiangiogenic agent (5–8). The molecular and cellular basis of this resistance is still poorly understood. HB demonstrates an important phenotypic diversity with three main histological subtypes presenting a spectrum of differentiation states (9) that is reflected at the transcriptomic level (2, 3, 9–13). HB is usually treated by cisplatin-based neo-adjuvant and adjuvant chemotherapy and surgical removal, leading to >80% 5-year survival (14). However, patients diagnosed with advanced unresectable tumours, chemo-resistant tumours or recurrent disease have a poorer prognosis. Chemotherapy-related toxicity is also a major concern for patients with HB (15).

For many years, our team has generated and characterized mouse models of liver carcinogenesis dependent on the β-catenin pathway. Using Cre-lox recombination and more recently CRISPR/Cas9 approaches in mice, we have demonstrated that oncogenic activation of the β-catenin pathway in hepatocytes following loss of *Apc* expression (APC^ΔHep^ model) or stabilizing mutations in *Ctnnb1* (βcat^ΔEx3^ model) is sufficient to promote cell transformation and tumour initiation (16, 17). APC^ΔHep^ and βcat^ΔEx3^ mice develop at equal risk HCC and HB-like tumours that closely recapitulate the histological and transcriptional features of human β-catenin-mutated HCC and mesenchymal HB, respectively (17). These mouse models have proven to be valuable tools for deepening our understanding of the mechanisms governing hepatocyte transformation under the control of the β-catenin pathway, as well as for testing new diagnostic and therapeutic approaches (18–20).

In recent years, three-dimensional (3D) *in vitro* culture systems have attracted considerable interest due to their potential to bridge the gap between two-dimensional (2D) culture systems and *in vivo* observations, particularly in the field of cancer (21–24). Among them, 3D tumouroids more accurately mimic the architecture and functions of the native tumour by maintaining cell-cell interaction, cell-extracellular matrix interaction, and by reproducing cellular heterogeneity and complexity of the microenvironment as closely as possible. Various methods for tumouroid establishment have been developed, ranging from the simplest to the most sophisticated, such as ultra-low attachment (ULA) plate cultures, scaffold-based systems, organ-on-chips and bioreactors, each with its own advantages and limitations (25).

In this study, we sought to generate tumouroids from our clinically relevant mouse models using an original device based on dynamic suspension culture in a rotating bioreactor (ClinoStar® system, Celvivo) that favours and maintains 3D structures in a static orbit by counterbalancing gravitational forces. This technology does not require the addition of a matrix or any other type of additive that could disrupt gene expression. In addition, this culture type eliminates cell contact with plastic, minimizes shear forces due to the gentle flow of media and promotes uniform gas and nutrient exchanges. It has previously been reported that culturing the human liver cell line HepG2/C3A in the ClinoStar® produces 3D structures that are highly relevant for studying various aspects of liver pathophysiology (26–29). After optimizing the culture conditions for mouse hepatocytes activated for the β-catenin pathway, we report that this technology allows the maintenance of tumouroids derived from both HCC and HB-like mouse tumours with phenotypical and transcriptional features perfectly matching the tumours of origins. Our new 3D preclinical models show great potential for advancing both basic and translational research on β-catenin–activated liver cancers, as demonstrated by the sensitivity of HCC-derived tumouroids to WNTinib.

## Materials and methods

### Mouse models and procedures

For the generation of APC^Δhep^ liver organoids, APC^lox/lox^*/TTR-Cre^ERT2^* mice (2-3-month-old, males) were injected intraperitoneally with 2 mg tamoxifen (MP Biomedicals) diluted in corn oil (16). High dose tamoxifen induces *Apc* deletion in around 90% of hepatocytes (**Fig.S1)**, and the subsequent activation of β-catenin-dependent pathways throughout the liver parenchyma. For the generation of APC^Δhep^ liver tumouroids, *APC*^lox/lox^ mice (2-3-month-old, males) were injected with an adenovirus encoding Cre recombinase (AdCre, 0.5 × 10^9^ particles), resulting in APC loss in a fraction of hepatocytes (30-40%), subsequent β-catenin activation, and emergence of liver tumours within 4-6 months (16). For the generation of βcat^Δex3^ liver tumouroids, C57BL/6J mice were injected with two adeno-associated virus serotype 8 (AAV8) encoding saCas9 and sgRNA against *Ctnnb1* to promote the specific loss of *Ctnnb1* exon 3 in a subset of hepatocytes as previously reported (17). In both models, tumour development was followed by 2D-ultrasounds under anesthesia with isoflurane inhalation, as previously described (18, 19). The maximum cumulative tumour diameter permitted by the ethics committees was 12 mm and was respected. For terminal tissue and blood collection, mice were anesthetized with ketamine (80 mg/kg) and xylazine (5 mg/kg). Mice were housed on a 12/12-h light/dark cycle at 22°C-24°C with *ad libitum* access to food (Safe A03, SAFE® Complete Care Competence) and water. All animal experiments were carried out in accordance with the French government regulations and with the approval of two ethical committees (Charles Darwin, Sorbonne Université, APAFIS#16420, APAFIS#45107 and CEEA34, Université de Paris, APAFIS#14472).

### ClinoReactor preparation

One day before their use for organoid or tumouroid culture, the humidification beads contained in the ClinoReactors (CelVivo) were hydrated with 20 ml sterile ultra-pure water (CelVivo). Then, the ClinoReactors were thoroughly washed with DPBS for 15 min before cell seeding at a rotation speed of 15 rpm as previously described (27).

### Generation of hepatic organoids from APC^Δhep^ mice

APC^Δhep^ mouse livers were perfused with collagenase six days following tamoxifen administration, and hepatocytes and non-parenchymal cells (NPC) were isolated and counted as previously described (30). Hepatocyte viability was systematically verified by trypan blue exclusion and was greater than 90%. For 2D culture, 500,000 hepatocytes were seeded on 6-well plates coated with rat tail collagen type 1 (Gibco) in William’s E medium with GlutaMAX™ (Gibco) supplemented with 10% fetal bovine serum (FBS), 100 U/mL penicillin/10 μg/mL streptomycin, 4 μg/mL insulin (Sigma-Aldrich), 0.1% bovine serum albumin (BSA) (Eurobio), 0.5 μg/mL amphotericin B (ThermoFisher Scientific), and 40 ng/mL dexamethasone (Sigma-Aldrich). Four hours later, the medium was removed and replaced with the same medium without FBS, insulin, BSA and amphotericin B. For 3D culture, hepatocytes and NPC were resuspended in William’s E medium with GlutaMAX™, supplemented with 10% FBS, 50 U/mL penicillin-streptomycin, and 1% insulin-transferrin-selenium (ITS-G, ThermoFisher Scientific). For culture in 96-well ULA plates (Greiner), a mixture of 2,000 hepatocytes together with 1,000 NPC was used per well. For dynamic culture in ClinoStar®, 500,000 hepatocytes and 250,000 NPC were seeded in a ClinoReactor rotating at 45 rpm. After four days, the medium from 3D cultures was removed and replaced with the same medium without FBS. The medium was then renewed every three to four days. All cultures were maintained at 37°C in a humidified atmosphere containing 5% CO_2_.

### Generation of hepatic tumouroids from APC^Δhep^ and βcat^Δex3^ mice

The tumours were excised and immediately placed on ice in basal medium DMEM F-12 (Gibco) until processing. The tumours were then minced gently into small pieces using fine scissors, transferred to a 50-mL centrifuge tube containing 10 mL of digestion solution (0.75 mg/mL collagenase (Sigma-Aldrich) with 0.15 U dispase (STEMCELL^TM^ technologies) in basal medium DMEM F-12) and incubated at 37°C for 20 min under gentle agitation. Three to four enzymatic digestion rounds were performed, depending on tumour size and type. After digestion, the cell mixture was filtered through a 100-µm cell strainer (Falcon) to remove aggregates and low-speed centrifuged (50 x g for 5 min at room temperature) to separate hepatocytes (pellet) from NPC (supernatant). NPC were centrifuged at 300 x g for 5 min. Hepatocytes and NPC were resuspended in 10 mL of HepatiCult™ Organoid Growth Medium (Mouse) (STEMCELL^TM^ technologies) and seeded in a ClinoReactor. The cell suspension was cultured in the ClinoStar® at 45 rpm for up to 16 days, with the medium replaced every 3-4 days.

### Organoid and tumouroid treatment with WNTinib

After 8 days of culture in the ClinoStar®, organoids and tumouroids were transferred into two ClinoReactors: one containing DMSO as a control condition and the other containing WNTinib (MedChemExpress) at 1 µM for organoids and 5 µM for tumouroids. The treatments were maintained for five days and renewed every two days.

### Total RNA isolation, reverse-transcription and quantitative real-time PCR (RT-QPCR)

Total RNA was extracted from liver tissues, organoids and tumouroids using TRIzol^TM^ reagent (ThermoFisher Scientific) according to the manufacturer’s protocol. cDNA was synthesized using Maxima reverse transcriptase (ThermoFisher Scientific). QPCR was performed with specific primers using BlasTaq^TM^ 2X PCR master mix (abm®) or TaqMan^TM^ gene expression assays (Applied Biosystems). The expression levels of each target gene were normalized to those of *B2m,* the most stable normalization gene under our conditions, and expressed as the relative quantity using the formula 2^−ΔΔCt^. For some experiments, a mixture of liver tissue cDNA served as a calibrator. PCR primer pairs and TaqMan gene expression assays are listed in **Table S1.**

### Histology, immunohistochemistry (IHC) and TUNEL assay

Hematoxylin-Eosin (HE) staining, IHC and TUNEL assay were carried out on formalin-fixed paraffin-embedded liver tissue sections as previously described (19). For IHC, liver sections were deparaffinized and hydrated, and antigen retrieval was performed in citrate buffer pH 6.0 for 20 min in a water bath at 95°C. The sections were blocked with TBS with 0.1% Tween (TBST) and 5% normal goat serum (Vector Laboratories), incubated with primary antibody for 1 h at room temperature or overnight at 4°C depending on the antibody (**Table S2**), washed twice with TBST and then incubated with biotinylated secondary antibody (1:750 and 1:200 dilutions for mouse and rabbit antibodies, respectively). Immunoreactivity was detected with an avidin biotin complex–3 (VECTASTAIN® Elite ABC-HRP kit, Peroxidase, Vector Laboratories), 3,3′-diaminobenzidine system (DAB substrate kit, peroxidase (HRP) with nickel, Vector Laboratories). Nuclei were then stained with Mayer’s hemalun solution, and the slides were mounted after dehydration. To perform IHC on organoids and tumouroids, samples were fixed for 1 h in 10% buffered formalin (VWR) at room temperature, washed with PBS, then covered with 2% agarose in a special mould (Leica) before being embedded in paraffin. Detection of *in situ* apoptosis was performed using the terminal deoxynucleotidyl transferase dUTP nick-end labeling (TUNEL) assay, following the manufacturer’s instructions (Click-iT™ TUNEL Alexa Fluor™ 647, ThermoFisher Scientific). Photographs of entire slides were captured using AxioScan Z1 scanner (Zeiss) for IHC slides, or with Eclipse E800 microscope (Nikon) and Axio Observer Z1 microscope (Zeiss) for fluorescent and TUNEL slides. The number of stained cells per field was quantified using QuPath software (31). The surface area of spheroids/tumouroids was quantified using ImageJ software.

### RNA sequencing (RNA-seq) analysis

RNA-seq was performed on total RNA extracted from APC^Δhep^ non-tumour tissues (n=5), tumours (n=5 for HCC; n=4 for HB), and tumouroids established from these tumours. Of note, two HCC (tumours 4.B and 4.G) were developed in the same mouse liver. Fastq files were then aligned using STAR algorithm (version 2.7.6a), on the Ensembl release 101 reference. Reads were then count using RSEM (v1.3.1) and the statistical analyses on the read counts were performed with R (version 3.6.3) and the DESeq2 package (DESeq2_1.26.0) to determine the proportion of differentially expressed genes between two conditions. The standard DESeq2 normalization method (DESeq2’s median of ratios with the DESeq function), was used with a pre-filter of reads and genes (reads uniquely mapped on the genome, or up to 10 different loci with a count adjustment, and genes with at least 10 reads in at least 3 different samples). Following package recommendations, the Wald test was used with the contrast function and the Benjamini-Hochberg FDR control procedure to identify the differentially expressed genes. R scripts and parameters are available at https://github.com/GENOM-IC-Cochin/RNA-Seq_analysis. Heatmaps were generated using the web-based tool Morpheus (https://software.broadinstitute.org/morpheus), and KEGG pathway enrichment analyses were performed using ShinyGO 0.85.1. Datasets are available at GSEXXX.

### Multiplexed immunofluorescence

Multiplexed immunofluorescence was performed on the Comet^TM^ instrument (Lunaphore) on 5-µm paraffin-embedded non-tumour, HCC, and tumouroid sections from APC^Δhep^ model in a 12×12 mm chamber. Slides underwent deparaffinization followed by heat-induced epitope retrieval in alkaline buffer pH 9.0 at temperature reaching 102°C, followed by controlled cooling prior to subsequent staining steps. Tissues were submitted to 16 cycles of antibody incubation/elution/image acquisition. One or two primary antibodies were added for 8 min and one or two adhoc secondary antibodies for 4 min. Used antibodies are described in **Table S2**. The fields of view were saved and the background was subtracted using the HORIZON software (Lunaphore).

### Statistical analysis

Differences between two sample groups were assessed using a Mann-Whitney test. For comparisons involving more than two groups, Kruskal-Wallis test was performed. A p-value < 0.05 was considered statistically significant.

## Results

### Generation of liver organoids from Apc^ΔHep^ mice using ClinoStar® system

For many years, 2D culture of human and murine hepatocytes on collagen matrix has been the gold standard method. However, the rapid cell dedifferentiation and cell mortality associated with this culture method have called it into question (32). Accordingly, 2D culture impacted identity of APC^KO^ hepatocytes isolated from *Apc^lox/lox^/TTR-Cre^ERT2^*mice injected with a high dose of tamoxifen (**Fig.1A**). Freshly isolated APC^KO^ hepatocytes overexpressed three major and direct β-catenin target genes, *Glul* encoding glutamine synthetase (GS) (100-fold increase), *Axin2* (100-fold increase) and *Lgr5* (30-fold increase) compared to wild-type (WT) hepatocytes isolated from *Apc^lox/lox^* mice injected with tamoxifen (**Fig.1B**). After 24 hours in 2D culture, *Glul*, *Axin2* and *Lgr5* were still overexpressed in APC^KO^ hepatocytes compared to WT hepatocytes, but their fold change was lower than that observed in freshly isolated APC^KO^ hepatocytes (20-, 50-, and 0.5-fold increase, respectively). Thus, the maintenance of β-catenin-activated hepatocyte identity is rapidly altered by 2D monoculture.

**Fig. 1.**
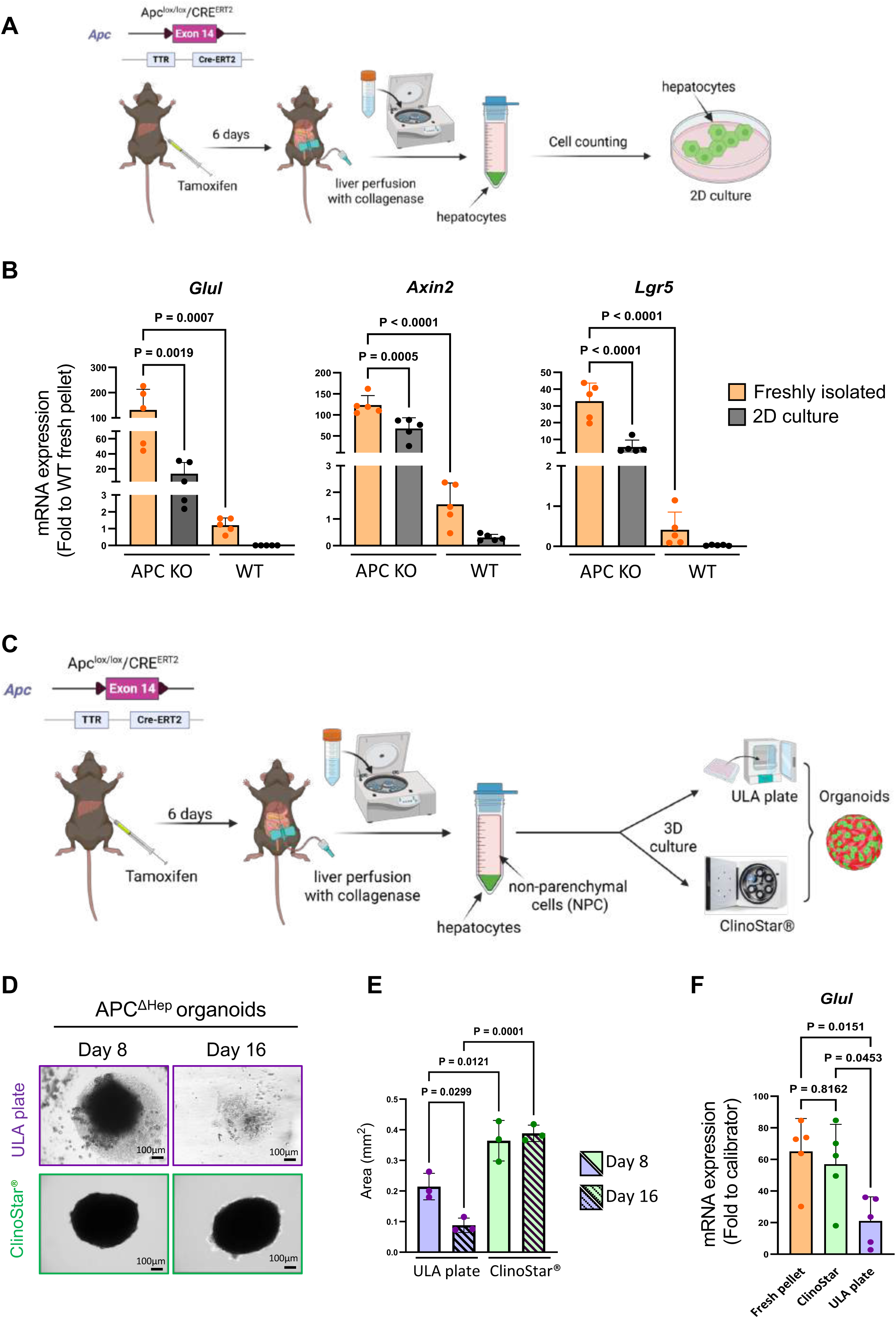
Generation of liver organoids from Apc^ΔHep^ mice in ULA plates and a ClinoStar®-based 3D culture system. **(A)** Schematic illustration of the experimental design for 2D culture of APC^KO^ hepatocytes (Created with BioRender.com). **(B)** Expression of *Glul*, *Axin2* and *Lgr5* transcripts assessed by RT-qPCR (n = 5) in freshly isolated APC^KO^ hepatocytes or after 24 h in 2D culture normalized to WT fresh pellets. **(C)** Schematic illustration of the experimental design for 3D culture of organoids (Created with BioRender.com) **(D)** Representative brightfield images of liver APC^ΔHep^ organoids acquired 8 or 16 days after cell plating in ULA plates or in the ClinoStar®. **(E)** Quantification of the organoid integrity area (mm^2^) at day 8 and 16 after plating in ULA plates or in the ClinoStar®. **(F)** Expression of *Glul* transcripts assessed by RT-qPCR (n = 5) under the different culture conditions. All bar graphs represent mean values ± SD, with individual dots corresponding to independent samples. For comparisons involving more than two groups, Kruskal-Wallis test was performed.

We therefore decided to switch to 3D culture of APC^KO^ hepatocytes together with NPC (immune, mesenchymal and endothelial cells) (**Fig.1C**). To this aim, a 2:1 ratio of Apc^KO^ hepatocytes/NPC were seeded in a ClinoReactor and cultured during eight days in a ClinoStar®. As a comparison, Apc^ΔHep^ organoids were also produced by static culture on ULA plates (**Fig.1C**). By preventing cell adhesion to the plastic dish, ULA plates promote the spontaneous aggregation of cells into spheroids (33). A major limitation of this technique is the formation of a necrotic core due to poor diffusion of nutrients and oxygen (34). After eight days of culture in the ClinoStar®, APC^KO^ hepatocytes mixed with NPC formed around 40 well-individualized organoids of uniform area per clinoreactor (**Fig.1D-E**). High inter-experimental reproducibility was obtained with the ClinoStar® system regarding organoid area and number. Organoids also formed systematically in each well of the ULA plates, but they were smaller than those generated with the ClinoStar® (**Fig.1D-E**). After 16 days in culture, 100% of liver organoids retained a compact, and rounded morphology in the ClinoStar®, unlike those cultured in ULA plates, 62% of which showed structural loosening and tended to be smaller (**Fig.1D, E**). These findings highlight the advantage of dynamic rotational culture in preserving organoid structural integrity and probably viability and proliferation features over extended periods.

At the molecular level, *Glul* mRNA expression was similar between Apc^ΔHep^ organoids generated in the ClinoStar® after eight days in culture and freshly isolated APC^KO^ hepatocytes, whereas *Glul* expression was significantly reduced in Apc^ΔHep^ organoids generated in ULA plates (**Fig.1F**). Histology and IHC analyses were then performed to better characterize Apc^ΔHep^ organoid structures obtained from both conditions (**Fig.2A**). After staining with HE, numerous cells lacking nucleus were observed in the core of organoids obtained from the ULA plates suggesting cell death, whereas such cellular entities were not observed in the organoids from the ClinoStar® (**Fig.2B**). Consistently, the detection of cleaved caspase-3 revealed strong staining in organoids generated with ULA plates but not with the ClinoStar® (**Fig.2B-C**), indicating that the ClinoStar® device limits cell death in Apc^ΔHep^ organoids. Apc^ΔHep^ organoids produced in the ClinoStar® also showed homogeneous GS labelling as well as membrane, cytoplasmic and nuclear staining of β-catenin. TUNEL assay revealed an increased number of positive cells in organoids generated in ULA plates compared to those generated in the ClinoStar®, confirming enhanced cell apoptosis in ULA plates (**Fig.2D**). Importantly, NPC were detected within Apc^ΔHep^ organoids. Thus, macrophages and endothelial cells were evidenced using F4/80 and CD146 immunostaining, respectively (**Fig.2B-C**). Finally, higher level of cell proliferation (KI67 staining) was observed in Apc^ΔHep^ organoids produced in the ClinoStar® compared to ULA plates (**Fig.2B-C**). The presence of dead cells in Apc^ΔHep^ organoids obtained using ULA plates had an impact on the quality of IHC, which was either difficult to detect (GS, cytoplasmic and nuclear β-catenin) or accompanied by significant background noise and non-specific staining (KI67, F4/80, CD146) (**Fig.2B**). Overall, these data indicate that the ClinoStar® system offers favourable conditions for the production of β-catenin-activated liver organoids by allowing the maintenance of different cell populations in the absence of cell death.

**Fig. 2.**
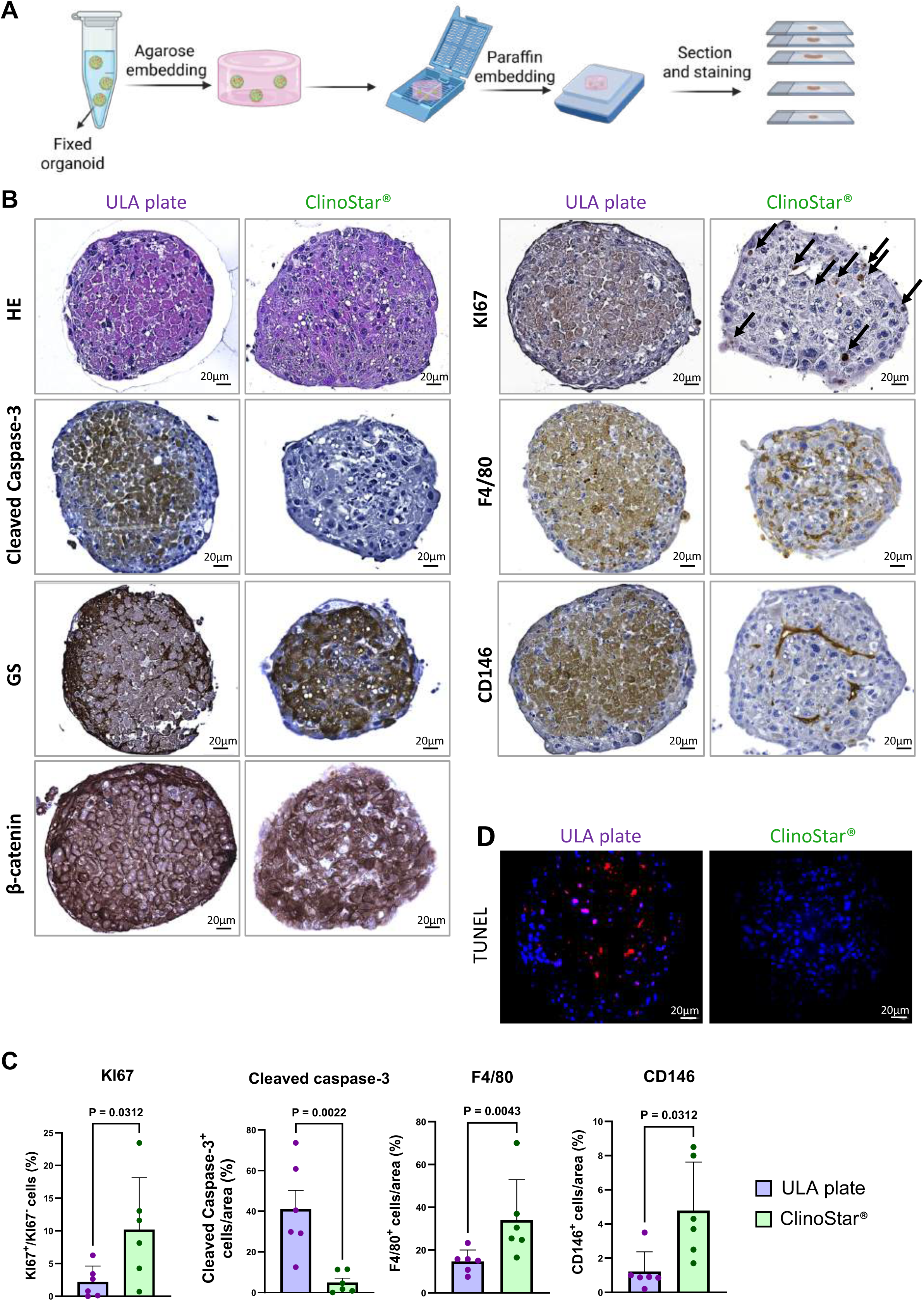
Beyond ULA plates: 3D culture in a ClinoStar® improves the development of APC^ΔHep^ liver organoids. **(A)** Schematic illustration of the spheroid inclusion protocol (Created with BioRender.com). **(B)** Representative HE staining and immunostaining for cleaved capase-3, GS, β-catenin, KI67, F4/80 and CD146 in APC^ΔHep^ organoids generated after 8 days in ULA plates (*left panel*) or in the ClinoStar® (*right panel*). The black arrows indicate nuclei positively stained for KI67. **(C)** Quantitative analysis of KI67, cleaved caspase-3, F4/80 and CD146 positive cells in organoids cultured in ULA plates or ClinoStar® (n = 6). **(D)** Representative images of apoptotic cells detected by TUNEL staining (red), with nuclei counterstained with DAPI (blue) in organoids cultured during 8 days in ULA plates (*left panel*) or in the ClinoStar® (*right panel*). All bar graphs represent mean values ± SD, with individual dots corresponding to independent samples. Differences between two sample groups were assessed using a Mann-Whitney test.

### Generation of liver tumouroids from Apc^ΔHep^ and βcat^Δex3^ mouse tumours using ClinoStar® system

These data led us to investigate whether ClinoStar® system could be also useful for generating β-catenin-activated tumouroids. To this end, *Apc*^lox/lox^ mice were injected with an adenovirus encoding Cre recombinase (AdCre) and followed by ultrasounds for tumour development during several months. Once tumours had been detected, they were harvested, dissociated mechanically and enzymatically, and seeded in bioreactors (**Fig.3A**). Two types of tumours were obtained from these mice, either well-differentiated APC^Δhep^ HCC (n=5) or undifferentiated APC^Δhep^ HB-like tumours (n=4).

**Fig. 3.**
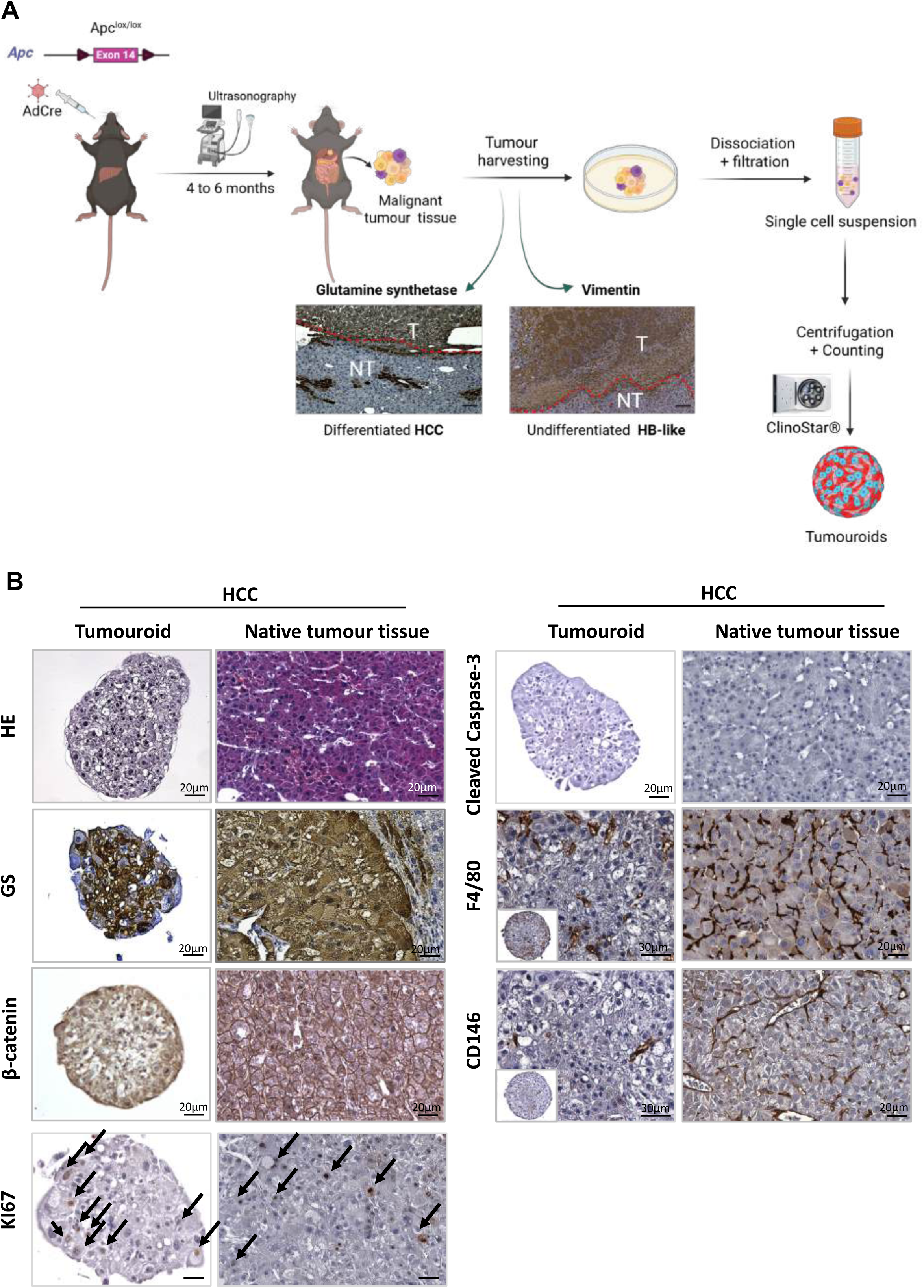
Culture in the ClinoStar® generates and preserves Apc^ΔHep^ HCC tumouroids. **(A)** Schematic illustration of the experimental design for 3D culture (Created with BioRender.com). **(B)** Representative HE staining and immunostaining for cleaved capase-3, GS, β-catenin, KI67, F4/80 and CD146 in tumouroids formed 8 days after plating in the ClinoStar®, compared to their native HCC. The black arrows indicate nuclei positively stained for KI67.

After eight days of culture in the ClinoStar®, we observed that tumour cells and NPC isolated from both HCC and HB-like tumours formed 3D structures, with approximately 30–40 tumouroids per culture and diameters ranging from 250 to 300 µm. These 3D structures reproduced the structural and cellular complexity of their native tumour tissue. Thus, APC^Δhep^ HCC tumouroids retained β-catenin pathway activation, evidenced by strong cytoplasmic staining of GS and membrane, cytoplasmic and nuclear staining of β-catenin (**Fig.3B**). As expected, HB-like tumouroids show less intense GS staining together with strong vimentin staining, underlying their mesenchymal status (**Fig.4**). No cell death (cleaved caspase-3 staining) was detected in the core of tumouroids, regardless of the type of the tumour of origin, while proliferative hepatocytes (KI67 staining) were detected throughout these 3D structures. As observed in the preneoplastic organoids (**Fig.2**), F4/80 and CD146 staining revealed the presence of macrophages and endothelial cells, respectively. Besides conventional IHC with chromogenic detection, tumouroids from APC^ΔHep^ HCC were also analyzed using a high-throughput automated multiplex immunofluorescence staining platform which relies on high-pressure microfluidic circuits. As shown in **Fig.S2**, the detection of proteins specific for the different cell subtypes was clearly evidenced within APC^Δhep^ HCC tumouroids.

**Fig. 4.**
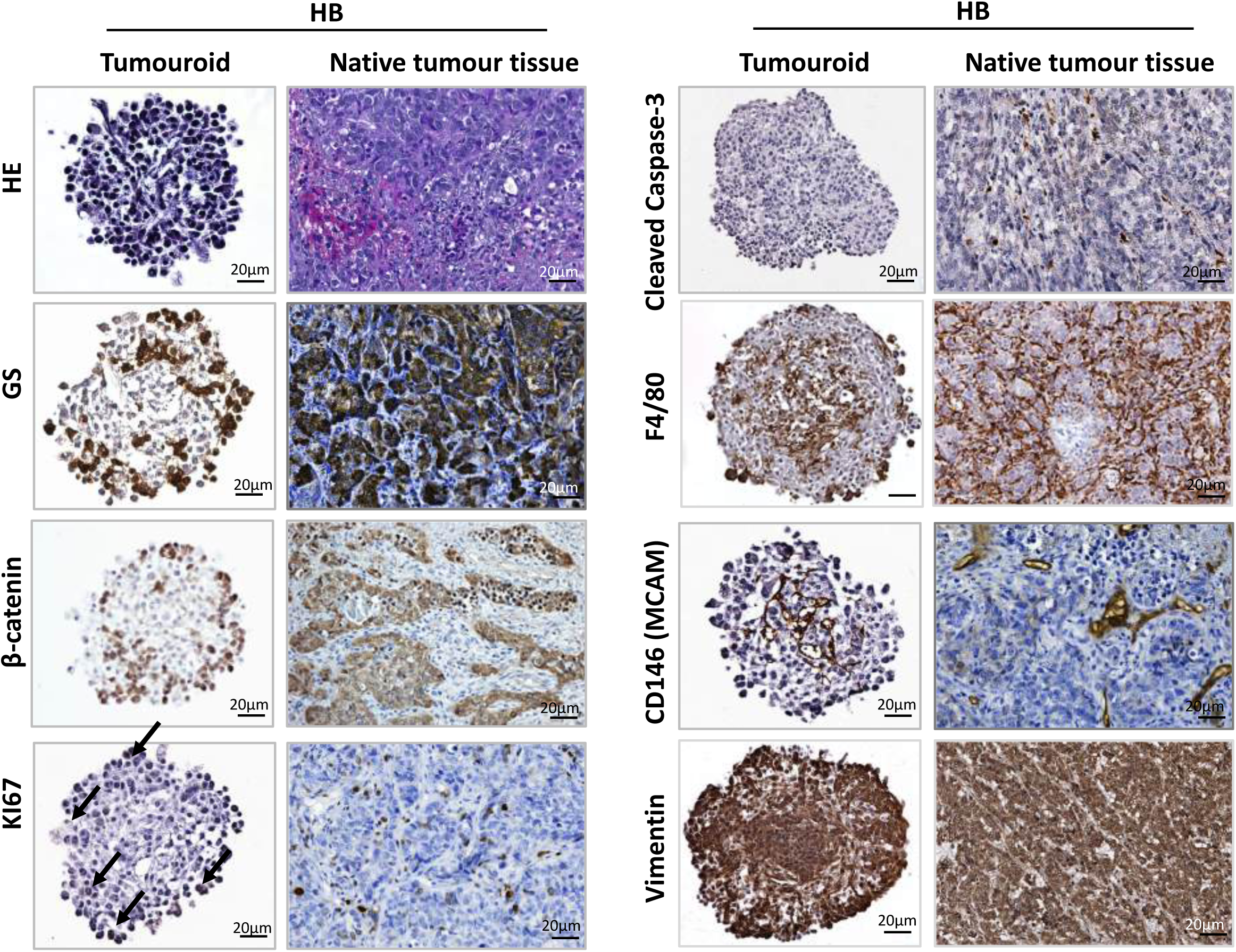
Culture in the ClinoStar® generates and preserves Apc^ΔHep^ HB-like tumouroids. Representative HE staining and immunostaining for cleaved capase-3, GS, β-catenin, KI67, F4/80 and CD146 in tumouroids formed 8 days after plating in the ClinoStar® compared to their native HB-like tumours. The black arrows indicate nuclei positively stained for KI67.

We confirmed these data with βcat^Δex3^ HCC which is equivalent to APC^ΔHep^ HCC at the transcriptomic level (17). Tumouroids derived from βcat^Δex3^ HCC retained the histological architecture from their original tumour (**Fig.S3**). Altogether, these data suggest that the dynamic culture in the ClinoStar® is efficient to generate tumouroids from both murine β-catenin-activated HCC and HB-like tumours, which preserve intra-tumoural cellular heterogeneity.

### APC^Δhep^ tumouroids cultured in the ClinoStar® maintain the transcriptome of the tumour of origin

We further performed RNA-seq to characterize APC^Δhep^ HCC (n=5) and HB tumouroids (n=4) compared to their native tumours and adjacent non-tumour tissues. Principal component analysis (PCA) indicated that each tumouroid was perfectly clustered with its tumour of origin, while non-tumour tissues were clearly separated into a tight cluster (**Fig.5A** for HCC and **Fig.6A** for HB). Samples from HCC#1/tumouroid#1 appeared to be outliers. The data for HCC#1 showed that it was a larger tumour (12 mm compared with 7-8 mm for the others), exhibiting signs of apoptosis (data not shown). Differentially expressed genes were then analyzed by MA plots, which also revealed tiny differences between each tumouroid and its tumour of origin (**Fig.5B** for HCC and **Fig.6B** for HB). Only *Fam177a2* and *2810047C21Rik* (a RIKEN-annotated mouse gene with unknown function) were significantly upregulated in HB-like tumouroids compared with tumours (**Fig.6B**). We then examined the expression pattern of genes specific to each type of tumouroids compared to non-tumour liver tissue (**Fig.5C** and **Fig.6C**; samples HCC#1/tumouroid#1 were excluded from this analysis). As previously described in APC^Δhep^ HCC (30, 35–37), HCC tumouroids expressed high levels of *Ctnnb1* targets genes (both canonical and liver specific genes), genes involved in glutamine, bile and drug metabolisms, inflammation, redox signaling as well as low levels of genes involved in amino acid catabolism (**Fig.5C**). Similar to the APC^Δhep^ HB-like tumours (17, 38), HB-like tumouroids expressed high levels of canonical *Ctnnb1* target genes as well as Apc^Δhep^ embryonic liver genes, mesenchymal markers and Hippo signature (**Fig.6C**). Transcripts encoding vascular markers such as *Mcam*/Cd146 were also significantly induced in both Apc^Δhep^ HCC and HB-like, confirming our observations in IHC (**Fig.3B** and **Fig.4**). KEGG pathway enrichment analysis was also performed between tumouroids and their non-tumoural related samples. In HCC-derived tumouroids, some of the pathways showing the highest enrichment scores were related to cell cycle and xenobiotic and gluthatione metabolisms (**Fig.5D**) while pathways related to amino acid metabolism were among the most downregulated (**Fig.5E**). Several pathways related to mitosis and cell division showed high enrichment scores in HB-derived tumouroids (**Fig.6D**) while several pathways related to lipid metabolism were downregulated (**Fig.6E**). This analysis confirms substantial pathway-level differences between tumouroids derived from HCC and HB and their non-tumoural tissues, which are consistent with the biological characteristics of the native tumours: a more metabolic phenotype for HCC and a more proliferative phenotype for HB-like tumours. Collectively, these data demonstrate that our protocol and culture conditions generate murine HCC and HB-like tumouroids which recapitulate the complexity and oncogenic programs of tumours of origins.

**Fig. 5.**
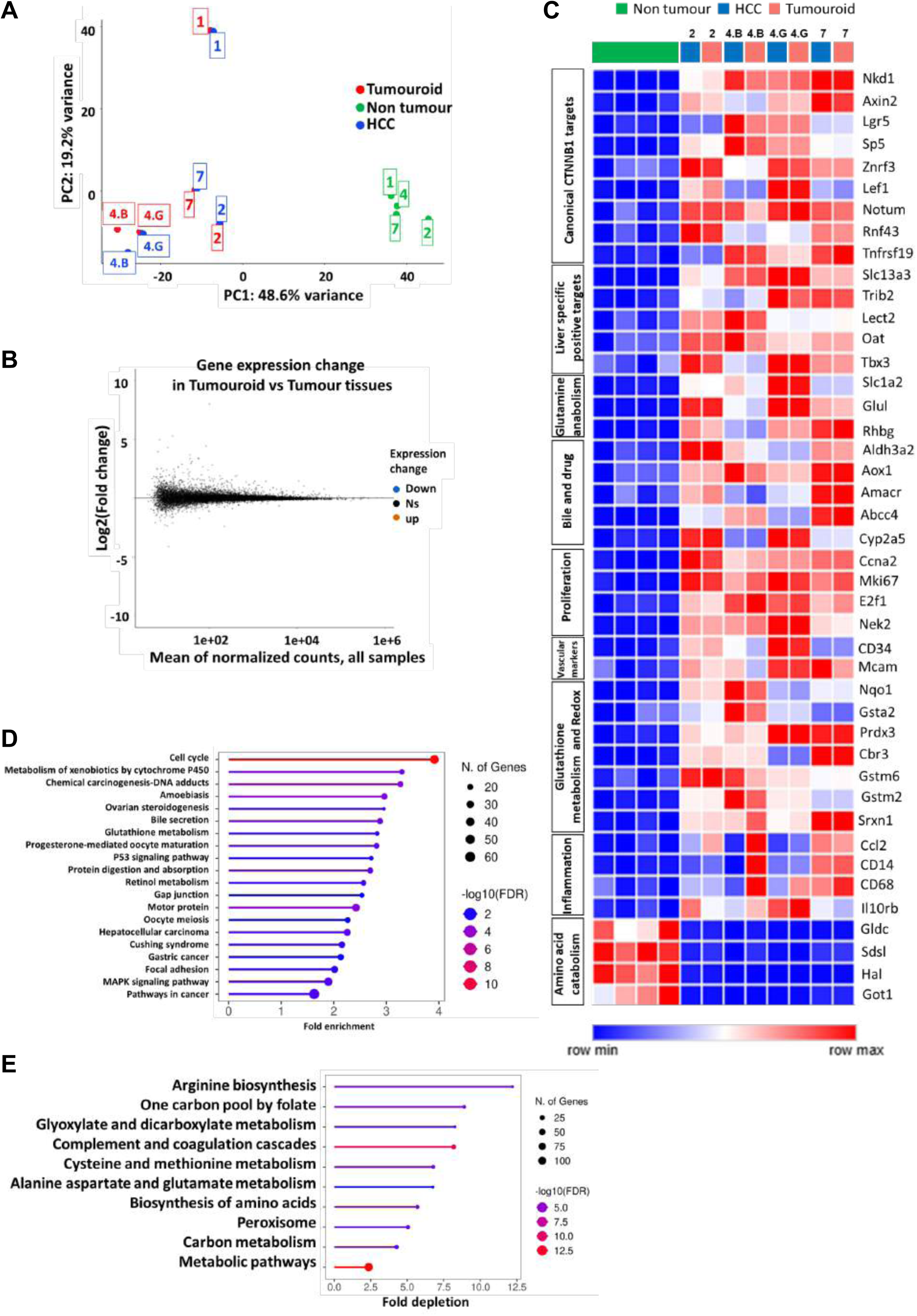
Culture in the ClinoStar® maintains the transcriptomic programs of Apc^ΔHep^ HCC tumouroids. **(A**) PCA of normalized gene expression profiles from tumouroids, HCC and adjacent non-tumoural livers. **(B)** MA plot showing gene expression between each tumouroid and its corresponding HCC. Each dot represents a gene, plotted as Log2 fold change versus the mean of normalized counts across all samples. Significantly upregulated genes are shown in orange, downregulated genes in blue, and non-significant genes in grey. **(C)** Heatmap of differentially expressed genes between tumouroids, HCC, and adjacent non-tumoural livers. Unsupervised hierarchical clustering was performed on samples. **(D)** KEGG pathway enrichment analysis of genes significantly upregulated in tumouroids compared to adjacent non-tumoural livers. **(E)** KEGG pathway enrichment analysis of genes significantly downregulated in tumouroids compared to adjacent non-tumoural livers.

**Fig. 6.**
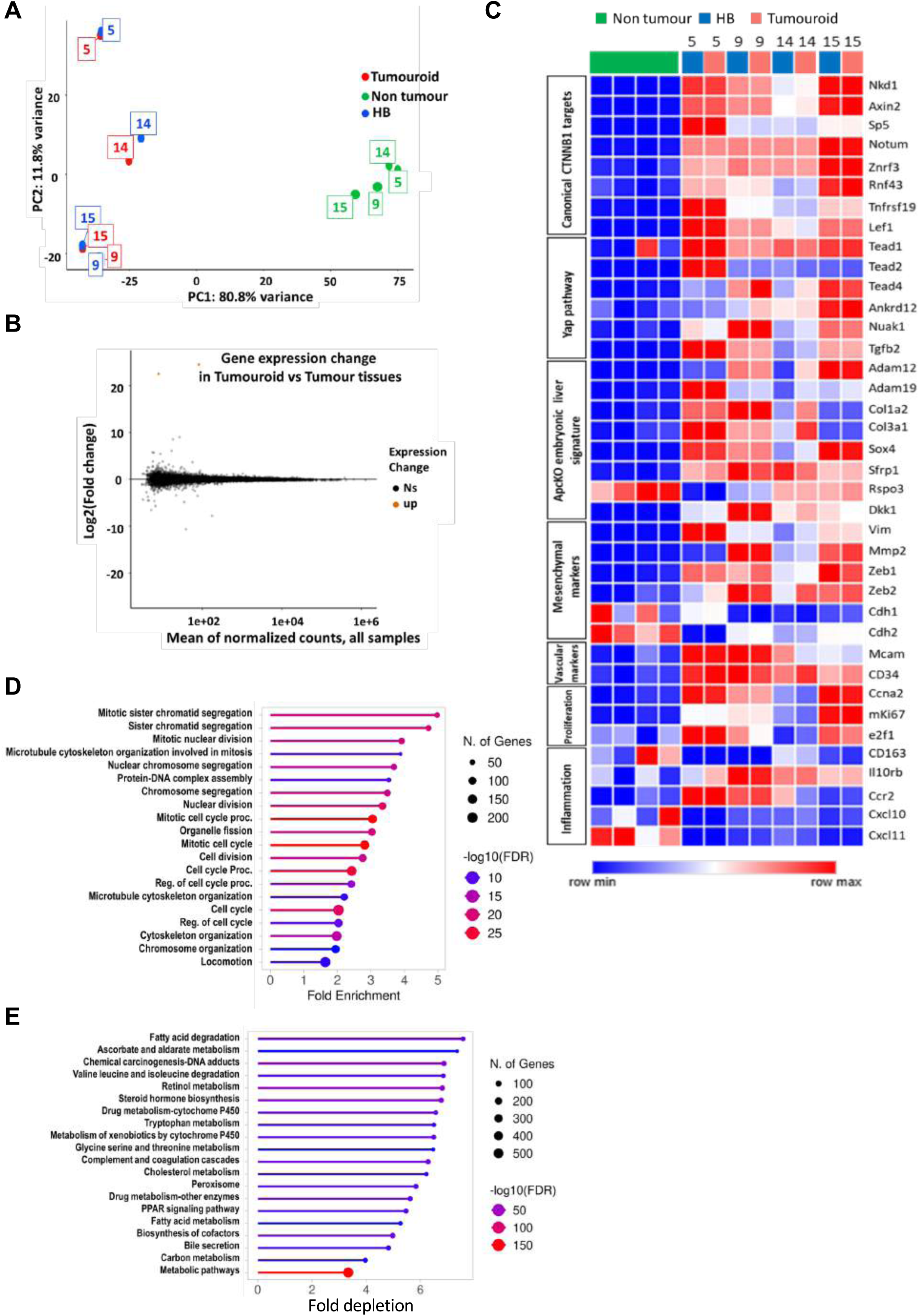
Culture in the ClinoStar® maintains the transcriptomic programs of Apc^ΔHep^ HB-like tumouroids. **(A)** PCA of normalized gene expression profiles from tumouroids, HB-like tumours and adjacent non-tumoural livers. **(B)** MA plot showing gene expression between each tumouroid and its corresponding HB-like tumour. Each dot represents a gene, plotted as Log2 fold change versus the mean of normalized counts across all samples. Significantly upregulated genes are shown in orange, downregulated genes in blue, and non-significant genes in grey. **(C)** Heatmap of differentially expressed genes between tumouroids, HB-like tumours and adjacent non-tumoural livers. Unsupervised hierarchical clustering was performed on both genes and samples. **(D)** KEGG pathway enrichment analysis of genes significantly upregulated in tumouroids compared with adjacent non-tumoural livers. **(E)** KEGG pathway enrichment analysis of genes significantly downregulated in tumouroids compared with adjacent non-tumoural livers.

### WNTinib suppresses proliferation and promotes apoptotic responses in β-catenin-activated organoids and tumouroids

To investigate whether organoids and tumouroids generated in the ClinoStar® could be new relevant preclinical tools for drug screening, we treated both APC^Δhep^ organoids and βcat^Δex3^ tumouroids with WNTinib, a recently identified multi-kinase inhibitor which demonstrated efficiency against human HCC and HB cells with *CTNNB1* activating mutations (39, 40). Once APC^Δhep^ organoids had formed after eight days of culture in the ClinoStar®, 1 μΜ WNTinib was added for five additional days. As observed in **Fig.7A** and **Fig.7B**, WNTinib induced marked structural and molecular changes. HE staining showed a disorganized structure with cells without nucleus within the core of the treated organoids compared to untreated ones. GS immunostaining was less intense under treatment with WNTinib, while cleaved caspase-3 was detected within treated organoids only. TUNEL assay revealed increased number of positive cells in WNTinib-treated organoids, confirming enhanced cell apoptosis (**Fig.S3,** *upper panel*). Proliferation was also markedly reduced in treated organoids as evidenced with KI67 immunostaining (**Fig.7A-B**). The repression of the β-catenin signalling pathway, associated with downregulation of *Glul* and *Axin2* mRNA as well as the induction of apoptosis, evidenced by the induction of the pro-apoptotic *Noxa* mRNA and the downregulation of anti-apoptotic *Bcl2l1* mRNA, were also demonstrated by RT-PCR in WNTinib-treated APC^Δhep^ organoids (**Fig.7C**). We also investigated whether βcat^Δex3^ HCC-derived tumouroids were sensitive to WNTinib. After eight days of culture, HCC tumouroids were treated with 5 μM WNTinib for five additional days in the ClinoStar®. WNTinib induced the appearance of cells without nucleus within tumouroid core which was associated with an apoptotic program revealed by increased caspase-3 cleavage (**Fig. 7D**) and TUNEL labelling (**Fig.S3,** *lower panel*) as well as increased *Noxa* and decreased *Bcl2l1* mRNA levels (**Fig.7F**). WNTinib also reduced the numbers of Ki67+ cells (**Fig.7D-E**). These data unveil both preneoplastic organoids and tumouroids as new powerful preclinical models for drug screening.

**Fig. 7.**
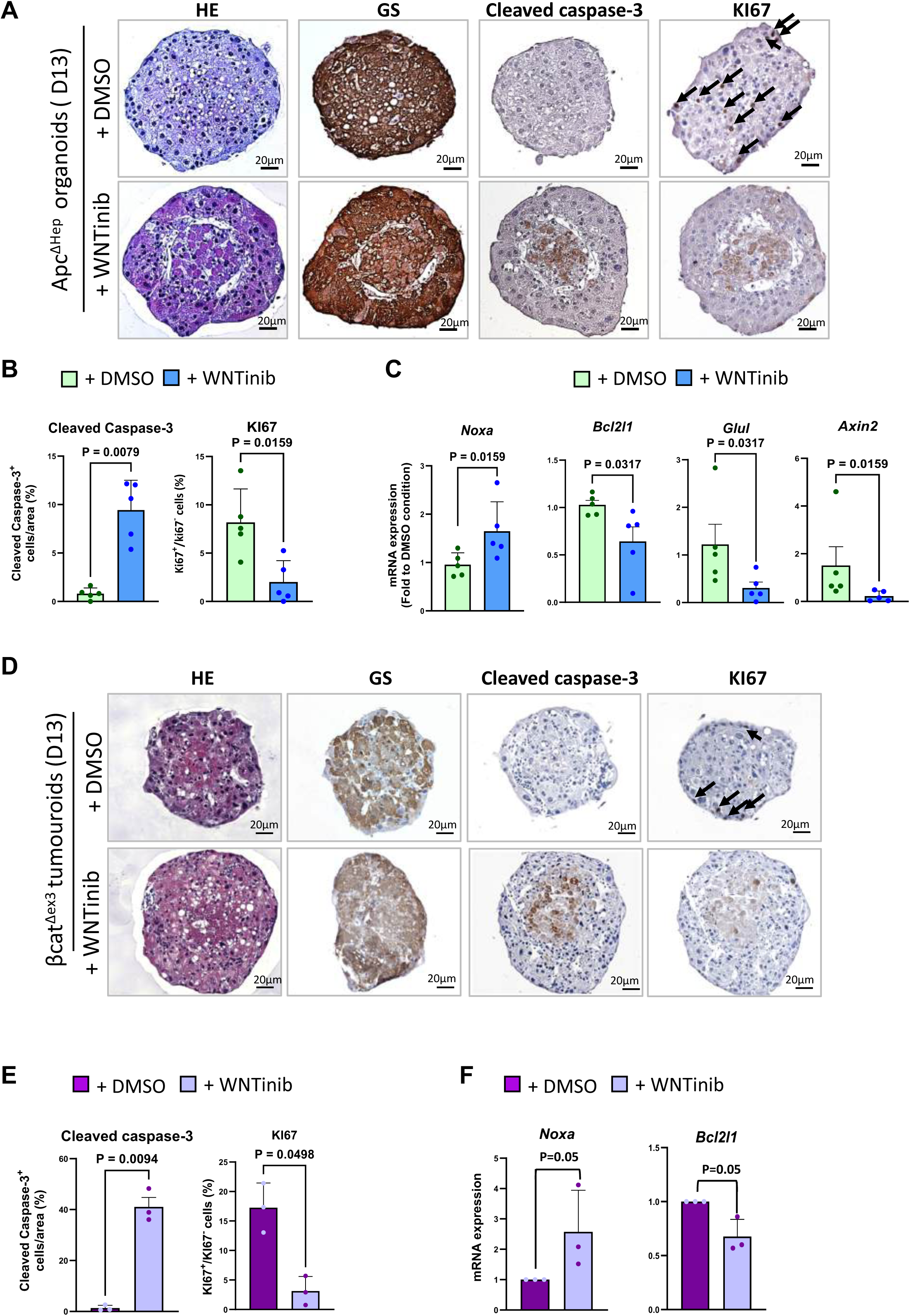
WNTinib induces apoptosis and impairs proliferation in Apc^ΔHep^ liver organoids and βcat^Δex3^ HCC tumouroids. **(A)** Representative HE staining and immunostaining for GS, cleaved caspase-3 and KI67 in Apc^ΔHep^ organoids formed 8 days after plating in the ClinoStar® and treated with WNTinib at 1 µM or DMSO for 5 days. The black arrows indicate nuclei positively stained for KI67. **(B)** Quantification of cleaved caspase-3 and KI67 positive cells in DMSO- and WNTinib-treated organoids (n = 5). **(C)** Expression of *Noxa*, *Bcl2l1*, *Glul* and *Axin2* transcripts evaluated by RT-qPCR (n = 5) in WNTinib- and DMSO-treated organoids. **(D)** Representative hematoxylin–eosin staining and immunostaining for GS, cleaved caspase-3 and KI67 in βcat^Δex3^ HCC tumouroids formed 8 days after plating in the ClinoStar® and treated with WNTinib at 5 µM or DMSO for 5 days. The black arrows indicate nuclei positively stained for KI67. **(E)** Quantification of cleaved caspase-3 and KI67 positive cells in DMSO- and WNTinib-treated βcat^Δex3^ HCC tumouroids (n = 3). **(F)** Expression of *Noxa* and *Bcl2l1* transcripts evaluated by RT-qPCR (n = 3) in DMSO- and WNTinib-treated βcat^Δex3^ tumouroids. All bar graphs represent mean values ± SD, with individual dots corresponding to independent samples. Differences between two sample groups were assessed using a Mann–Whitney test.

## Discussion

In the present study, we successfully established murine preneoplastic organoids from livers with hepatospecific β-catenin overactivation as well as tumouroids from β-catenin-activated well differentiated HCC and poorly differentiated HB-like tumours. IHC and gene expression analyses reveal that these organoids and tumouroids retain their histological architecture and the expression patterns of their tumour they derived from. This success is based on the use of a vertical rotary bioreactor which utilizes gentle rotational dynamics to achieve near-microgravity conditions, thus reducing shear stress while maintaining optimal nutrient and oxygen distribution (26). The morphological homogeneity and size reproducibility of liver organoids and tumouroids cultured in the ClinoStar® system are consistent with the findings of recent studies using this technology to establish brain organoids (41) and lung tumouroids (42). Moreover, organoids and tumouroids generated from mouse livers and tumours with β-catenin activation, respectively, showed very low levels of cleaved caspase-3, particularly in their core, indicating that dynamic culture limits hypoxia and the subsequent apoptotic cell death. The protocol we have developed is relatively simple, relies on commercially available cell culture media, and does not require a preliminary step of aggregating dissociated cells in hydrogel or micropattern plates, as has been previously reported for other tissues and cell types (43). Furthermore, this protocol does not involve the addition of extracellular matrix, which increases reproducibility across experiments due to the absence of undefined factors associated with matrix-based products. Interestingly, this culture method allows hepatic organoids and tumouroids to maintain cellular diversity for at least 8 days, as highly differentiated and undifferentiated tumour hepatocytes, macrophages and endothelial cells were detected within these structures. This shows that this type of culture helps preserve a certain degree of tumour complexity. While the ClinoStar® device has been described previously as capable of generating spheroids from the HepG2 liver cancer cell line (28, 44), our study is the first to report its use for the formation of organoids and tumouroids from preneoplastic hepatocytes and liver tumours. The therapeutic management of primary liver tumours, particularly tumours activated for the β-catenin pathway, still requires further research. Thus, HCCs with a β-catenin mutated pathway, which account for up to 40% of HCCs, exhibit immune exclusion (with regard to T lymphocyte infiltration) and, when inoperable, are more refractory to standard treatments that combine immunotherapies (8, 45–47). The treatment for HB consists of cisplatin-based chemotherapy before and after surgery (14). However, resistance to chemotherapy is observed in metastatic or recurrent cases and the toxicity of chemotherapy can be a major concern in young patients (14). It is therefore essential to identify other vulnerabilities, overcome chemoresistance and develop therapies with low toxicities to improve HCC and HB patient outcomes.

2D and 3D cultures of human HCC and HB cell lines have long been used as a reference for *in vitro* drug screening, but due to their clonal and single-cell type nature, these models cannot recapitulate tumour heterogeneity, which may limit data translational value. Patient-derived xenografts have proved to be useful models for human HCC and HB (23, 48, 49) because they retain histology and gene expression from their tumour of origin. However, the establishment of mouse cohorts with PDXs is time-consuming, not always successful (the engraftment efficiency may be low), expensive, and not suitable for large-scale drug sensitivity testing because it does not fit within the framework of human animal research. Therefore, tumouroid cultures represent an alternative resource for drug discovery to liver cancer PDXs. The Meritxell Huch group pioneered this field by developing matrix-embedded tumouroids from human HCC and cholangiocarcinoma (48). They showed that their tumouroids are reliable *in vitro* cancer models, recapitulating the main features of the different subtypes of HCC and cholangiocarcinoma analyzed. Since then, this technology has been used by other groups to develop tumouroids from different mouse models of liver carcinogenesis (44, 49).

Within this framework, our study provides proof of concept that dynamic culture in a ClinoStar® represents a suitable approach for generating tumouroids applicable to HCC and HB modeling and drug screening. From a small fragment of tumour, we were able to produce dozens of tumouroids from murine HCC and HB that are highly similar in terms of shape, size, and cellular content and mirror the tumour of origin. The mouse models used in our study have demonstrated their clinical relevance in several studies. Thus, Apc^ΔHep^ mice with activation of the β-catenin pathway throughout the hepatic lobules, from which the spheroids described herein were derived, have proven useful for characterizing the early stages of constitutive activation of the β-catenin pathway in hepatocytes (24). Apc^ΔHep^ and βcat^Δex3^ mice exhibiting focal activation of the β-catenin pathway in the liver develop differentiated HCC and HB-like mesenchymal tumours that mimic human tumours at the histological and transcriptional levels (26). Murine HCC also exhibit metabolic traits such as glutamine anabolism and choline addiction (18), as well as responses to oxydative stress that mimic those observed in human HCC (35). Our new models should significantly reduce the number of mice used in pre-selection and evaluation tests for new therapeutic agents. In order to prove the concept that the murine tumouroids we generated from HCC would be useful for drug screening, we tested their sensitivity to WNTinib. WNTinib is a multikinase inhibitor which has been recently identified as a transcriptional repressor of mutant β-catenin and it has been evaluated in human liver cancer cell lines cultured in 2D and mouse- and patient-derived HCC and HB tumouroids cultured in Matrigel (39, 40, 44). In our dynamic culture, WNTinib potently inhibited proliferation and induced apoptosis in murine HCC tumouroids, consistent with its previously identified modes of action. This opens the way to test other drugs in our HCC and HB-like models.

In conclusion, we provide a new methodology for the generation of murine tumouroids from β-catenin-driven HCC and HB. These tumouroids represent valuable resources which should be useful tools for both basic cancer research and the development of precision medicine due to their translational and ethical advantages, especially their potential to reduce animal experimentation. Beyond these models, this methodology could be extended to tumours derived from other murine models of liver cancer, thereby broadening its applicability for studying diverse oncogenic contexts. Moreover, the compatibility of this approach with the generation of tumouroids from human liver tumour fragments is currently being investigated, which could further enhance its translational potential.

## Authors contributions

C.D.-M. and V.B.M. conceived the study and supervised all aspects of the project. V.B.M., F.L. and A.G. performed hepatocyte isolation and cell culture experiments. V.B.M. and A.H. performed histology, immunodetection and RT-qPCR. I.L. performed liver ultrasonography. SB followed tumour progression by ultrasonography. A.G. managed Apc^lox/lox^/TTR-Cre^ERT2^ mouse breeding. C.D.-M. and S.C. obtained funding supports. C.D.-M. and V.B.M. wrote the manuscript with contributions from all authors. All authors read and approved the final manuscript.

## Acknowledgements

We are grateful to CRC core facilities (INSERM UMRS_1138, Sorbonne Université, Université Paris Cité) and their team members for animal care and breeding (CFE, Mrs Sonia Prince), genotyping (CGB, Mrs Hermine Kakanakou), and cell imaging (CHICS, Dr Christophe Klein). We warmly thank the Institut Cochin’s core facilities (INSERM U1016, CNRS UMR8104, Université Paris Cité) and their team members for animal care and breeding (Moust’IC, Mr Marcio Do-Cruzeiro), liver ultrasonography (PIV, Dr Gilles Renault) and transcriptomic analysis (Genom’IC, Dr Franck Letourneur). Dr Véronique Blouin (CAPACITES, INSERM UMR1089, Université de Nantes, France) is thanked for recombinant AAV8 production. Dr Ronan Le Bouffant and Dr Helle Sedighi Frandsen are thanked for critical reading of the manuscript.

## List of abbreviations

AAV8: adeno-associated virus serotype 8
AdCre: adenovirus encoding Cre recombinase
BSA: bovine serum albumin
FBS: fetal bovine serum
GS: glutamine synthetase
HB: hepatoblastoma
HE: hematoxylin and eosin
IHC: immunohistochemistry
NPC: non-parenchymal cells
PCA: principal component analysis
PDX: patient-derived xenograft
2D: two dimensional
3D: three dimensional
ULA: ultra-low attachment
WT: wild-type

## Legends to supplementary figures

**Fig. S1.**
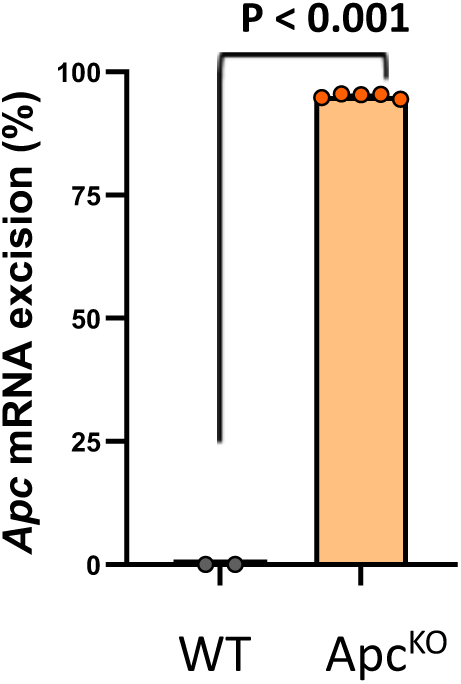
Expression of excised *Apc* mRNA. Percentage of *Apc* mRNA excision in freshly isolated WT and Apc^ΔHep^ hepatocytes evaluated by RT-qPCR (n = 5). Bar graphs represent mean values ± SD, with individual dots corresponding to independent samples. Differences between two sample groups were assessed using a Mann–Whitney test.

**Fig. S2.**
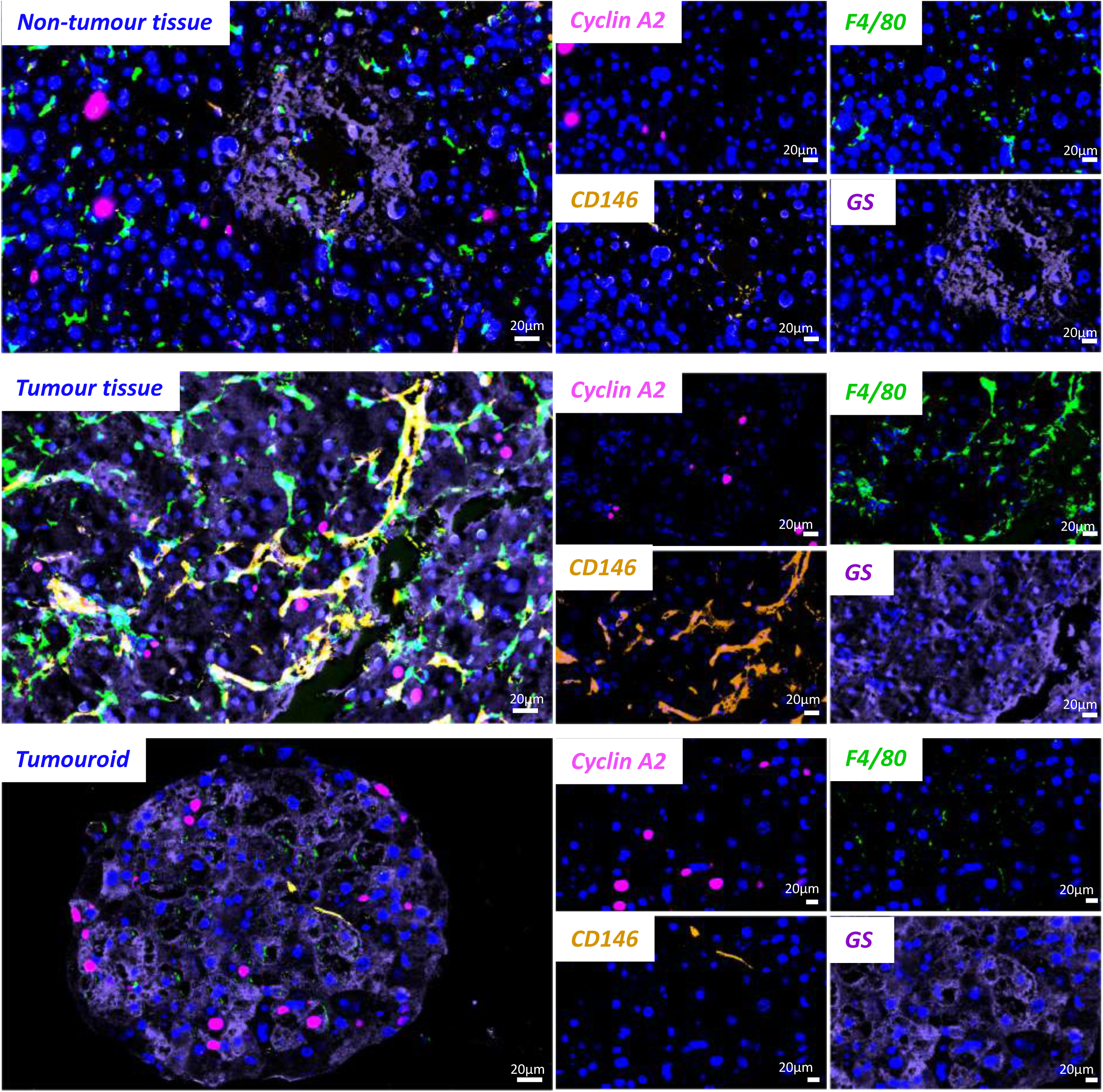
Multiplex tissue profiling of HCC-derived tumouroids. Multiplex immunofluorescence was performed using COMET^TM^ in non-tumour, HCC and tumouroids from Apc^ΔHep^ liver mouse. Representative COMET images showing cyclin A2 (magenta), CD146 (orange), GS (purple), F4/80 (green) and the nuclei counterstained with DAPI (blue).

**Fig. S3.**
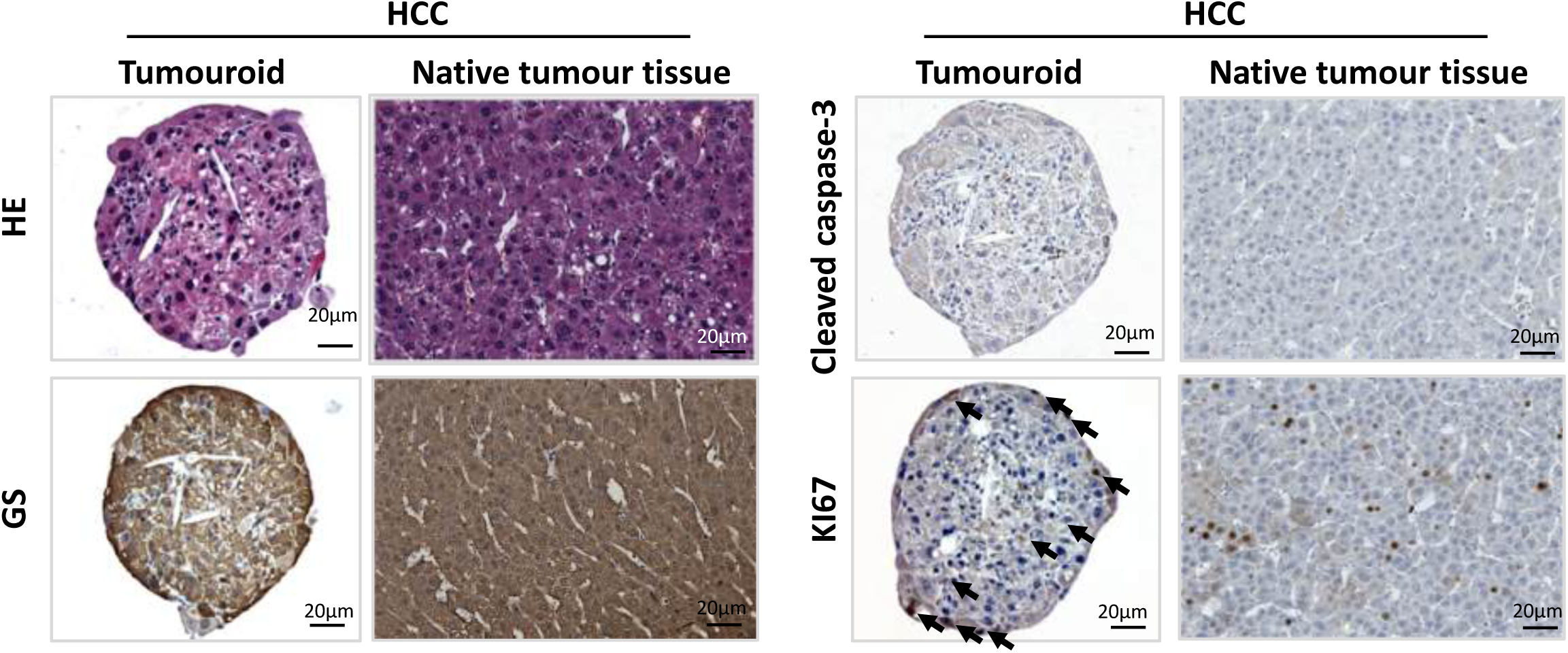
Histological and IHC characterization of βcat^Δex3^ tumouroids. Representative sections of HE staining and immunostaining for cleaved capase-3, GS, β-catenin, KI67, F4/80 and CD146 in β-cat^Δex3^ tumouroids formed 8 days after plating in the ClinoStar®. The black arrows indicate nuclei positively stained for KI67.

**Fig. S4.**
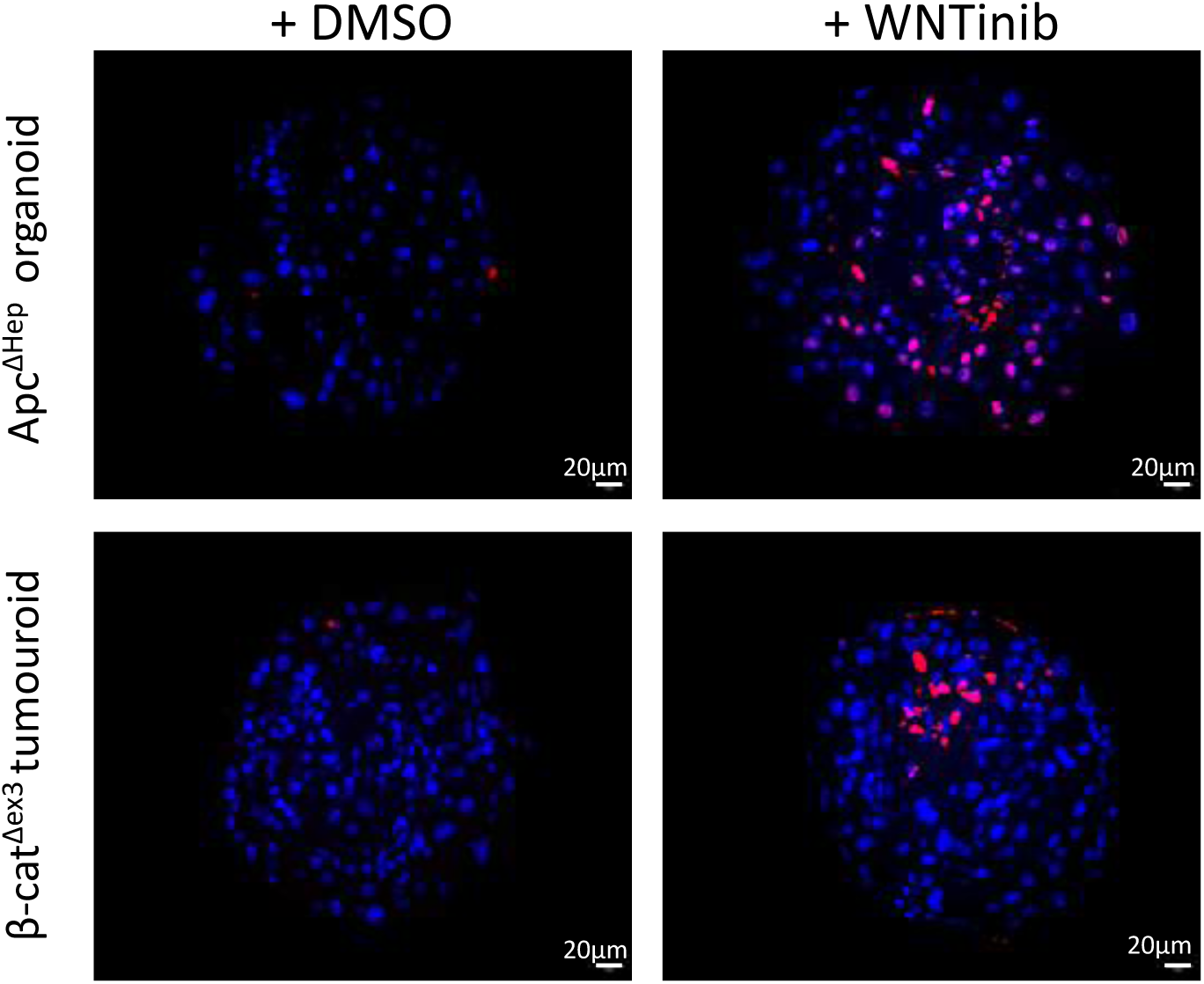
TUNEL assay in Apc^ΔHep^ liver organoids and βcat^Δex3^ HCC tumouroids. Representative images of Apc^ΔHep^ liver organoids (*top row*) and βcat^Δex3^ HCC tumouroids (*bottom row*) cultured during 8 days in the ClinoStar® system and treated with WNTinib at 1 µM (organoids) or 5 µM (tumouroids) or with DMSO for 5 days. Apoptotic cells were detected by TUNEL staining (red), with nuclei counterstained with DAPI (blue).

## Supplementary tables

**Table S1:**
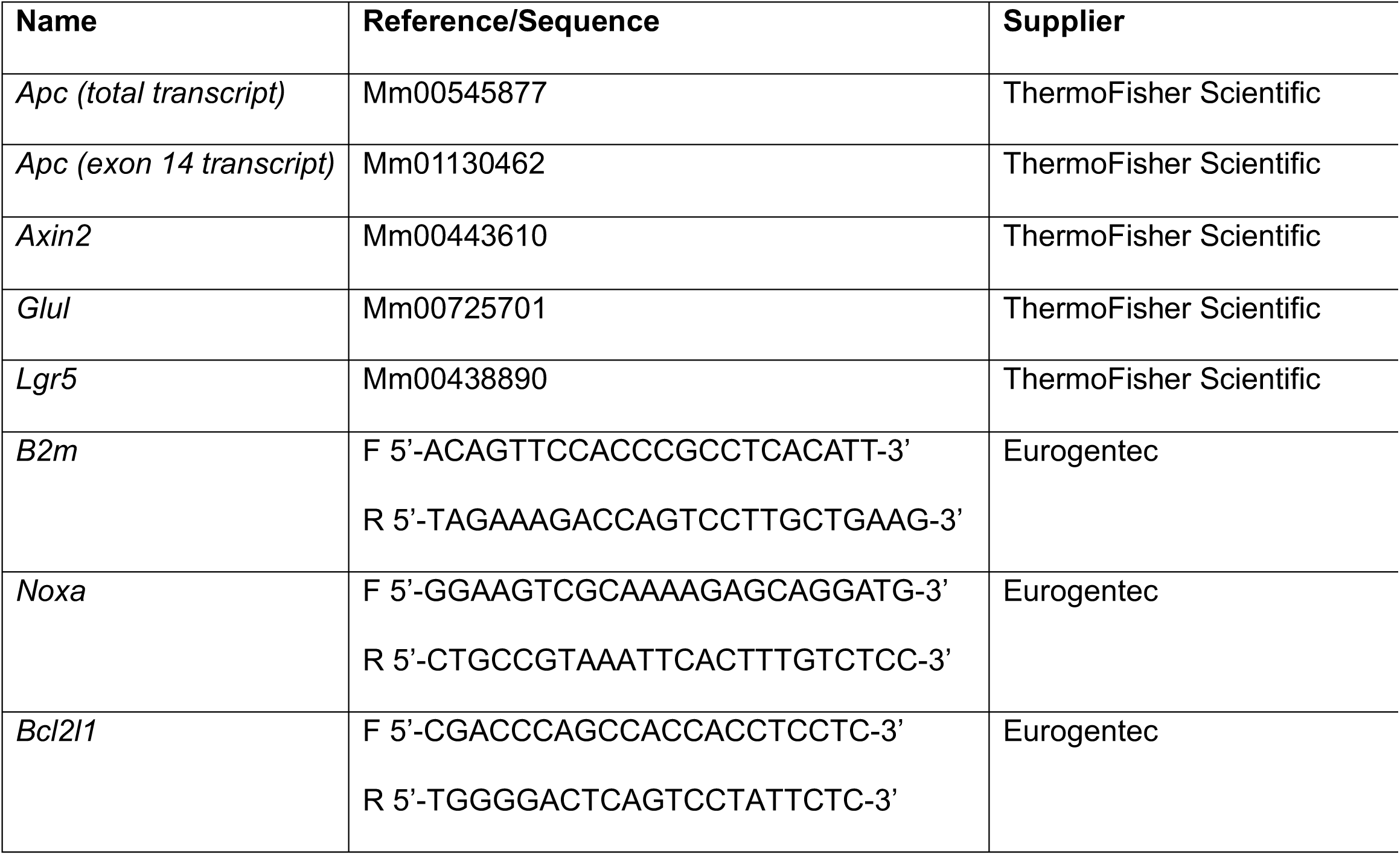
List of primers and probes.

**Table S2:** List of antibodies used in IHC and multiplex staining.

| **Name** | **Supplier** | **Reference** | **Dilution** | **Application** |
| --- | --- | --- | --- | --- |
| Cyclin A2 | Abcam | AB-32386 | 1:100 | Comet |
| CD146 | Cell Signaling | 81701 | 1:50, 1:100 | Comet / IHC |
| F4/80 | Cell Signaling | 70076 | 1:100 | Comet / IHC |
| β-catenin | Abcam | AB-224803 | 1:200 | Comet |
| Vimentin | Abcam | AB-92547 | 1:1000 | Comet |
| Glutamine synthetase | BD Biosciences | 610518 | 1:2000, 1:1000 | Comet / IHC |
| Anti-Rabbit AF555 | ThermoFisher Scientific | A31572 | 1:100 | Comet |
| Anti-Rabbit AF647 | ThermoFisher Scientific | A21245 | 1:200 | Comet |
| Anti-Mouse AF647 | ThermoFisher Scientific | A21236 | 1:200 | Comet |
| Anti-Rat AF647 | ThermoFisher Scientific | A21247 | 1:200 | Comet |
| Anti-Goat 647 | ThermoFisher Scientific | A21446 | 1:200 | Comet |
| KI67 | ThermoFisher Scientific | MA5-14520 | 1:200 | IHC |
| Cleaved caspase-3 | Cell signaling | D175 | 1:600 | IHC |

